# Pseudo-craniotomy of a whole-brain model reveals tumor-induced alterations to neuronal dynamics in glioma patients

**DOI:** 10.1101/2023.12.22.573027

**Authors:** Christoffer G. Alexandersen, Linda Douw, Mona L.M. Zimmermann, Christian Bick, Alain Goriely

## Abstract

Brain tumors can induce pathological changes in neuronal dynamics both on a local and global level. Here, we use a whole-brain modeling approach to investigate these pathological alterations in neuronal activity. By fitting a Hopf whole-brain model to empirical functional connectivity, we demonstrate that phase correlations are largely determined by the ratio of interregional coupling strength and intraregional excitability. Furthermore, we observe considerable differences in interregional-versus-intraregional dynamics between glioma patients and healthy controls, both on an individual and population-based level. In particular, we show that local tumor pathology induces shifts in the global brain dynamics by promoting the contribution of interregional interactions. Our approach demonstrates that whole-brain models provide valuable insights for understanding glioma-associated alterations in functional connectivity.

## 1 Introduction

One of the ultimate goals of neuroscience is to understand and control neuronal dynamics across scales, from the individual neuron to the whole brain. At the larger end of the scale, researchers have focused on imaging methods such as functional magnetic resonance imaging (fMRI), electroencephalography (EEG), and magnetoencephalography (MEG) to measure whole-brain dynamics. These methods have revealed consistent correlation patterns in neuronal activity across brain regions. Interestingly, these correlation patterns, commonly known as functional connectivity, are disrupted in diseases such as Alzheimer’s disease [1], depression [2], schizophrenia [3], and glioma [4–6]. Following these observations, computational neuroscientists have been formulating *whole-brain models* to explain functional connectivity patterns and how they change during disease. However, even in the simplest whole-brain models and for simple brain states, fitting the model output to functional connectivity is a significant challenge. Overcoming this challenge is crucial, as it is a major roadblock for using whole-brain models in clinical applications which has the potential to improve diagnosis, to understand disease mechanisms, and to enable personalized treatments [7–9]. For example, patients with brain tumors such as gliomas exhibit pathological alterations in functional connectivity that are not yet fully understood; the development of accurate whole-brain models can help reveal the underlying mechanisms of tumor pathology. Here, we employ a whole-brain model to elucidate the mechanisms that lead to functional connectivity alterations in glioma patients and determine the impact tumor regions have on the emergent whole-brain dynamics.

It is now well established that glioma patients have been shown to exhibit seemingly pathological differences in functional connectivity patterns across imaging modalities when compared to healthy controls [4–6]. In particular, overall functional connectivity has been shown to be lower in the default-mode network as measured by fMRI [10–13]. Similarly, lower functional connectivity is observed in MEG for higher frequency bands [14–17], whereas lower frequency bands quite consistently display higher functional connectivity compared to controls instead [15, 17, 18]. Further differences in functional connectivity have been observed in network metrics of MEG/fMRI functional connectivity such as clustering and path length [15, 18–20]. However, it remains unclear why and exactly how such connectivity deviations occur in some patients, but not all [21].

Whole-brain models are formulated to simulate empirical functional connectivity patterns and have been used recently to study a variety of diseases. Essentially, a whole-brain model is a network of coupled oscillators in which each oscillator produces a time series mimicking the oscillatory signals of a region found through neuroimaging modalities. These simulated signals can then be used to compute simulated functional connectivity which is then compared to experimental functional connectivity. The basic network architecture of a whole-brain model comes from structural reconstructions of the human brain, called structural connectomes. In a structural connectome, the network’s nodes denote brain regions of various sizes and the network’s edges encode some measure of physical connectivity between the regions; the most common measure being the integrity of axonal bundles divided by their length as reconstructed from diffusion magnetic resonance imaging (dMRI). The dynamical model of neuronal activity at each node varies from simple phenomenological oscillators to high-dimensional biophysical models. Whole-brain models have long been used to replicate resting-state functional connectivity as measured by fMRI, EEG, and MEG [22–24]. Additionally, whole-brain models have been used to investigate brain diseases and neuropsychiatric disorders, such as epilepsy [25, 26], Alzheimer’s disease [27–30], schizophrenia [31–33], and glioma [34–37].

Whole-brain models can be used to simulate the effect of lesions to better understand the effect of surgical craniotomy, which often is necessary for epilepsy and glioma patients [34]. In such studies, a set of brain regions is removed from the physical brain network structure of the whole-brain model (virtual surgery), and the resulting changes in functional connectivity are analyzed. The effect of brain region removal on the emergent functional connectivity patterns is nontrivial and depends on the structural connectivity network [35, 38]. For example, the effect of removing certain regions on fMRI-derived functional connectivity depends largely on the region. The removal of some regions—such as those in the cortical midline and the temporoparietal junction—results in global changes, whereas other regions—such as primary sensory and motor regions—result only in local changes [34]. Another highlight is the resilience of brain dynamics to surgical removal and the importance of node network measures in determining the impact of lesions [35]. Furthermore, by optimizing the parameters of the whole-brain model to glioma patients and control subjects, differences in neuronal dynamics can be inferred. For example, using fMRI-derived functional connectivity and a reduced Wong-Wang model for neuronal dynamics, local inhibitory connections were considerably lower in tumor regions whereas no significant difference was found in long-range, interregional connections [36]. Moreover, optimal parameter fits of the (reduced Wong-Wang) whole-brain model remain relatively stable after comparing before and after real-world craniotomy, and the capability of whole-brain models to accurately predict the effects of craniotomy in glioma patients is, as of yet, difficult to ascertain due to the small cohort sizes of glioma patients currently available [37]. Furthermore, considering changes induced by glioma in EEG/MEG, a whole-brain model approach demonstrated that tumor-induced changes in local neuronal dynamics can considerably reduce the overall functional connectivity strength, suggesting that local pathology can lead to global symptoms [39].

In this study, we use a whole-brain model of Hopf oscillators to explain the functional connectivity differences in glioma patients as compared to healthy controls, measured by MEG. We do so by identifying the optimal parameterization of the whole-brain model to empirical phase-lag index connectivity computed from MEG data of glioma patients and healthy controls. We find that the goodness of fit of the Hopf whole-brain model is invariant under a particular scaling law for the parameters. This scaling law demonstrates that the phase dynamics is almost solely determined by the *normalized coupling strength* which we define as the ratio of interregional coupling and intraregional excitability. Furthermore, we find that the optimal normalized coupling strength is higher in the glioma cohort than in the controls both on an individual and population-based level. We also show that the tumor regions generally induce increases in normalized coupling strength though there is considerable variability between patients. In summary, we find that the ratio of interregional and intraregional dynamics uniquely determines the phase dynamics of the Hopf whole-brain model and that tumors increase the relative contribution of interregional dynamics in glioma patients.

## 2 Methods

### 2.1 Participants

A total of *S* = 10 patients seen between 2011–2023 at Amsterdam UMC with suspected diffuse glioma were randomly selected from an ongoing prospective study on brain networks. Patients underwent MEG when glioma was suspected based on clinical history and MRI before tumor treatment or surgery was performed. Exclusion criteria were (1) age *<* 18 years, (2) psychiatric disease, (3) comorbidities of the central nervous system, (4) insufficient mastery of the Dutch language, and (5) inability to communicate adequately. After resection, molecular characteristics were assessed as part of the clinical routine, including prognostically favorable isocitrate dehydrogenase (IDH) mutations and 1p/19q codeletions [40]. This led to three subgroups: IDH-wildtype glioma (glioblastoma), IDH-mutant, non-codeleted glioma, and IDH-mutant, 1p/19q-codeleted glioma. Additionally, *S* = 33 healthy controls were included, which are further described in [41]. We refer to these two cohorts as the *glioma cohort* and the *control cohort*. The VUmc Medical Ethical Committee approved this study, which was conducted following the principles of the Declaration of Helsinki. All participants provided written informed consent before participation.

### 2.2 MEG data acquisition and processing

MEG was recorded for 5min in supine position during eyes closed no-task resting-state in a magnetically shielded room (VacuumSchmelze GmBh, Hanau, Germany), using a 306-channel (102 magnetometers, 204 gradiometers) whole-head MEG system (Elekta Neuromag Oy, Helsinki, Finland) and a sampling frequency of 1250Hz. Anti-aliasing (410Hz) and high-pass filters (0.1Hz) were applied online. Preprocessing involved visual inspection, noisy channel removal, and noise removal in the remaining signals. Anatomical MRI was used for co-registration with the digitized scalp surface and the Automated Anatomical Labeling atlas [42] for parcellation of the cortical ribbon into 78 regions. Broadband time series of neuronal activity were then reconstructed for each region’s centroid [43] using a scalar beamformer approach [44]. Fast Fourier transforms filtered the time series into six frequency bands: delta (0.5-4Hz), theta (4-8Hz), lower alpha (8-10Hz), upper alpha (10-13Hz), beta (13-30Hz), and gamma (30-48Hz).

Median peak frequencies of the spectral density were also computed for the control cohort (for the parameterization of the whole-brain model). For this purpose, epochs were curated visually by discarding epochs without clear alpha frequency peaks in occipital brain regions. Then, the peak frequency of each region of interest in each epoch was computed by finding the frequency above 4Hz with the highest spectral density and discarding peaks that were less than twice the average spectral density of the background spectrum. The median peak frequency of each brain region was then computed across all subjects and epochs.

### 2.3 MRI data and structural connectome reconstruction

Control MRI data were obtained using a 3T MRI system (Philips Ingenia CX) with a 32-channel receive-only head coil at the Spinoza Centre for Neuroimaging in Amsterdam, The Netherlands. A high-resolution 3D T1-weighted image was collected with a magnetization-prepared rapid acquisition with gradient echo (MPRAGE; TR = 8.1 ms, TE = 3.7 ms, flip angle = 8°, voxel dimensions = 1 mm^3^ isotropic). This anatomical scan was registered to MNI space through linear registration with nearest-neighbor interpolation and was used for co-registration and normalization of other modalities (dMRI and MEG) to the same space.

Diffusion MRI was collected in the controls only, with diffusion weightings of *b* = 1, 000 and 2,000 s/mm2 applied in 29 and 59 directions, respectively, along with 9 nondiffusion weighted (*b* = 0s/mm2) volumes using a multiband sequence (MultiBand SENSE factor = 2, TR = 4.7 s, TE = 95 ms, flip angle = 90°, voxel dimensions = 2 mm3 isotropic, no interslice gap). In addition, two scans with opposite phase encoding directions were collected for blip-up blip-down distortion correction using FSL topup [45]. Structural connectomes were constructed by performing probabilistic anatomically-constrained tractography (ACT) [46] in MRtrix3 [47]. A tissue response function was estimated from the preprocessed and bias field corrected dMRI data using the multishell multitissue five-tissue-type algorithm (msmt 5tt). Subsequently, the fiber orientation distribution for each voxel was determined by performing multishell multitissue-constrained spherical deconvolution (MSMT-CSD) [48]. ACT was performed by randomly seeding 100 million fibers within the white matter to construct a tractogram, and spherical-deconvolution informed filtering of tractograms (SIFT, SIFT2 method in MRtrix3) [49] was then performed to improve the accuracy of the reconstructed streamlines and reduce false positives. For every participant, their respective 3D T1-weighted image was used to parcellate the brain into the 78 cortical regions. We then used this parcellation to convert the tractogram to a structural network, where weighted edges represented the sum of all streamlines leading to and from all voxels within two brain regions. For patients, tumor masks were manually drawn on a combination of T1 weighted MRI with and without contrast, and FLAIR. Then, the atlas regions overlapping with each patient’s tumor mask were considered tumor regions.

### 2.4 Computation of the phase-lag index as a measure of functional connectivity

In this study, we used the *phase-lag index* (PLI) as a measure of functional connectivity [50]. PLI is a measure of the average asymmetry in pair-wise phase differences of oscillatory MEG signals from different brain regions and ranges from 0 (symmetry) to 1 (asymmetry) for each pair of regions.

The phase-lag index is computed in the same way for each cohort. For each cohort, we have *S* subjects. For each subject *S* ∈{ 1, …, *S*}, we have *E*_*s*_ epochs, each consisting of a *N* ×*M* matrix, where *N* is the number of brain regions and *M* is the number of time points. We denote each matrix by 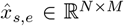 with dimensions *N* and *M*, where *e* denotes the epoch and *s* denotes the subject.

For processing, we apply a Butterworth filter to bandpass the signal into the alpha frequency band at 8–12 Hz. Thus, the Butterworth filter is a function *F* : ℝ ^*N ×M*^→ ℝ^*N ×M*^ that can be applied to each epoch. After bandpassing the data, we find the angle of its Hilbert transform Θ : ℝ^*N ×M*^ → ℝ^*N ×M*^. Next, we compute the phase-lag index connectivity. That is, we apply a function *P* : ℝ^*N ×M*^ → ℝ^*N ×N*^ to the data to construct a *N* -by-*N*, symmetric, functional connectivity matrix. This phase-lag index transformation is given explicitly by

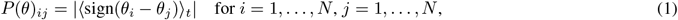

where ⟨ ⟩_*t*_ is the time average. We apply this process to compute the PLI for each matrix 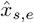:

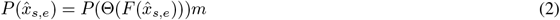

and compute the average experimental phase-lag index matrix for the entire cohort as

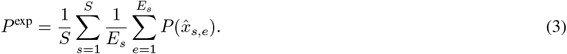

The experimental PLI matrices are noisy and are thus thresholded to ease the comparison with the noiseless whole-brain model. Thresholding the PLI matrix at a certain percentage *X*% means setting all elements in the lower *X*% percent to zero as ordered per magnitude. In some cases, we perform median thresholding instead, where all elements of the PLI matrix lower than the median of said matrix are set to zero.

### 2.5 The Hopf whole-brain model

We use a whole-brain model consisting of a network of Hopf oscillators. More specifically, we use the Hopf normal form (equivalent to Stuart-Landau oscillators) on a reconstruction of the physical human brain network, where the oscillators interact through a sigmoidal function without delays. As such, each Hopf oscillator denotes a brain region, where the links of the network denote axonal bundles connecting a pair of brain regions.

The human structural connectome network is represented by the adjacency matrix *W* with entries *w*_*ij*_. The state of the Hopf oscillators, *z*_*i*_ ∈ ℂ, are complex variables having real and complex parts *z*_*i*_ = *x*_*i*_ + i*y*_*i*_, where 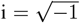. The Hopf oscillators are coupled through their real parts *x*_*i*_ and subsequent analysis of simulations will only consider the real part and not the imaginary part. Indeed, the real part *x*_*i*_ represents the simulated MEG signals while the imaginary part *y*_*i*_ can be thought of as a hidden variable needed to produce the necessary oscillatory dynamics.

The Hopf whole-brain model for a network *W* with *N* regions evolves through the following system of complex ordinary differential equations

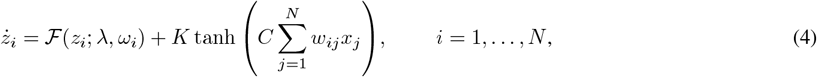

with

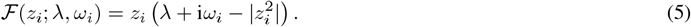

Here, *K* ∈ ℝ ^+^ = {*r* ∈ ℝ | *r*≥ 0} is the *global coupling strength, λ* R is the *global excitability parameter* (also known as the Hopf bifurcation parameter), *ω*_*i*_ ∈ ℝ ^+^ is the local natural frequency of oscillation, and *C* ∈ ℝ ^+^ is the *global scaling* of the structural connectome.

Each natural frequency *ω*_*i*_ is parameterized from the healthy control MEG data using the frequency peaks found for each brain region (see Section 2.2). Thus, the remaining free parameters are the coupling strength *K*, the excitability *λ*, and the structural connectivity scaling *C*.

### 2.6 Fitting whole-brain model to experimental functional connectivity

To assess the goodness of fit of the Hopf whole-brain model to experimental MEG data, we compare the functional connectivity of the simulated and experimental signals. The objective (cost) function we use to assess the goodness-of-fit is the Pearson correlation of the time-averaged pairwise phase-lag indices. We will now describe the settings of whole-brain model simulations, the processing of the simulated data, and the computation of the objective function.

We simulate the Hopf whole-brain model for 14.5 seconds and discard the first second of simulations to account for transient dynamics induced by the initial conditions. We save the simulated data with a sampling frequency of 1250 Hz. As such, the simulated data has the same length and sampling frequency as the experimental MEG data. The simulations are performed with the RK45 method (using the JiTCODE module for Python) with an absolute tolerance of 10^−6^ and a relative tolerance of 10^−3^.

As mentioned, we interpret the real part of the Hopf oscillator variables *x*_*i*_ as MEG signal and process it in the same manner as the experimental MEG signal. That is, we bandpass the signal into the alpha frequency band (8-12Hz) using a Butterworth filter (denoted by the function *F*) and subsequently apply the Hilbert transform to find the angles of the oscillators (denoted by the function Θ). We then compute the pairwise time-averaged phase-lag index of the processed simulated signal and compute the Pearson correlation of the simulated PLI matrix and the experimental PLI matrix.

Let *x* ∈ ℝ ^*N ×M*^ denote the simulated time series of *N* regions over *M* time points. Note that *x* is a function of the parameters and the initial conditions *z*(0) = *z*_0_ of the whole-brain model. As such, the simulated phase-lag index is *P* ^sim^ = *P* (Θ(*F* (*x*(*K, λ, C, x*_0_))), with *P*, *F*, and Θ defined above. Remembering that *P* ^exp^ denotes the experimental MEG PLI matrix averaged over epochs and subjects, the objective function is

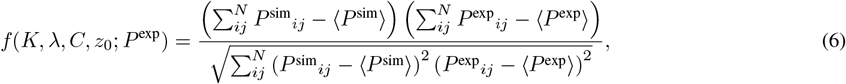

where ⟨⟩ denotes the average over all matrix entries. As the objective function depends on the initial conditions *z*_0_, we average the objective function over several runs with random initial conditions. For random initial conditions, each oscillator’s initial state *z*_*i*_ for *i* = 1, …, *N* is a random point in the unit disk |*z*_*i*_|≤ 1 picked with uniform probability.

In Section 3.3, we investigate the impact of tumor regions on the dynamics of the whole-brain model in individual patients and clusters of tumor regions. We do so by performing a *pseudo-craniotomy*, in which we remove the rows and columns corresponding to the tumor brain regions from both *P* ^sim^ and *P* ^exp^. In effect, the objective function ignores the tumor regions resulting in a change in optimal model parameters.

## 3 Results

### 3.1 Phase correlations emerge from the competition between inter- and intra-regional dynamics

To assess the goodness of fit for the Hopf whole-brain model, we compare simulations to experimental data. Specifically, we compute the phase-lag index of the simulated signals of the whole-brain model (example shown in Fig. 1a) and compare them with the averaged experimental phase-lag index of a healthy control group (experimental PLI found in Fig. 1c). The goodness-of-fit is defined as the Pearson correlation between the simulated and experimental pairwise phase-lag indices.

**Figure 1:**
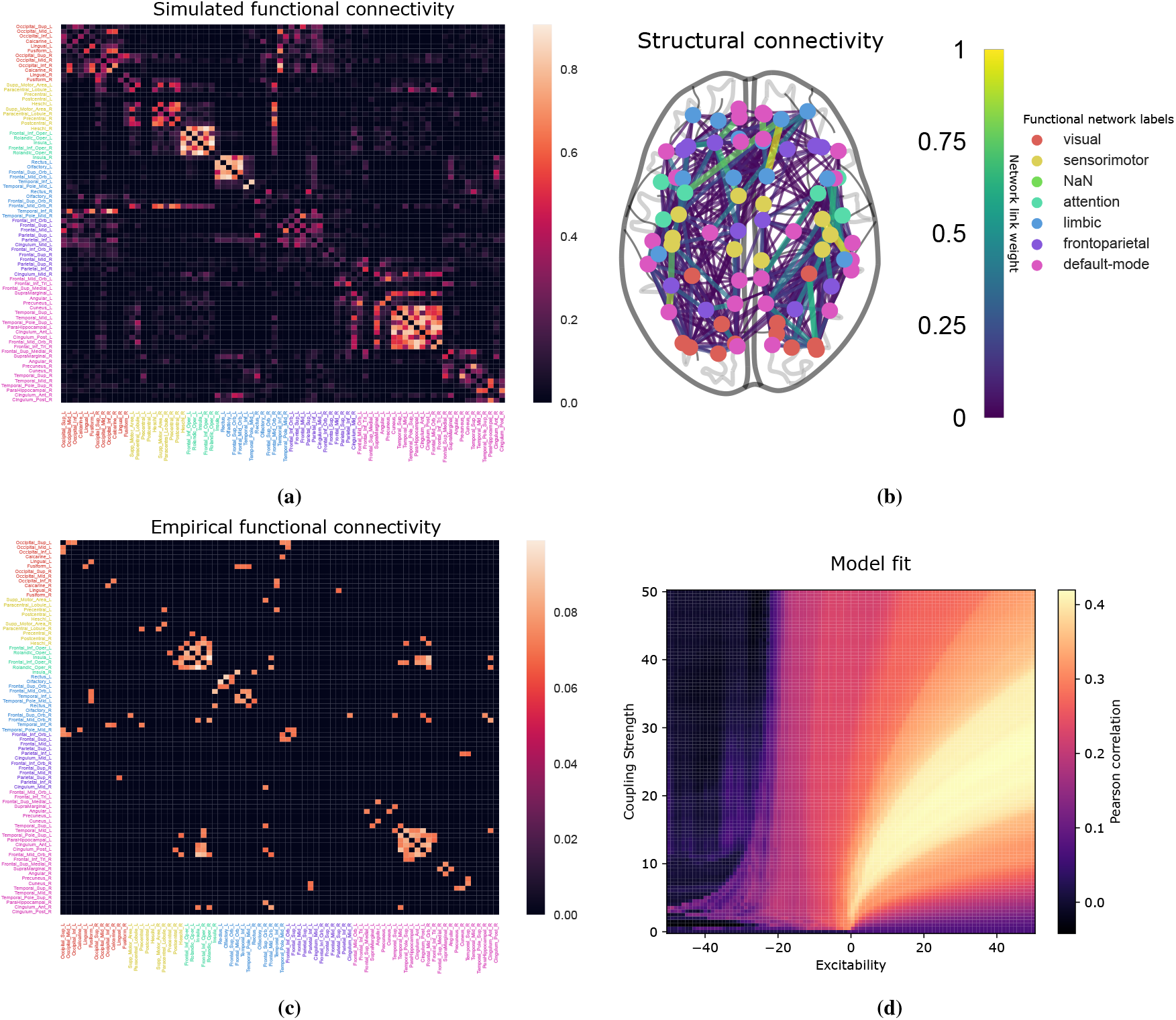
(a,c) An example of synthetic (simulated) phase-lag index from a Hopf whole-brain model and experimental PLI averaged over 33 healthy subjects from the control group. The name of each brain region is labeled and colored according to canonical functional networks (see legend to the upper right). (b) The weighted physical connectome averaged over the control group. Each node is colored similarly to the functional connectivity matrices. (d) The goodness of fit of the Hopf whole-brain model with respect to the coupling strength and excitability parameter. Each pixel shows the Pearson correlation between the simulated and experimental (control group average) phase-lag indices averaged over 500 whole-brain simulations with random initial conditions.

The experimental PLI is noisy and is thus thresholded to make it comparable to the noiseless, simulated PLI. Figure 10 in the Appendix shows the effect of thresholding on the average control PLI. We threshold the experimental, averaged PLI at the optimal threshold providing the best fit to the simulated data (see Fig. 11 in the Appendix). However, we note that the following results hold for a wide range of threshold values, as demonstrated in the Appendix. In particular, the results have also been reproduced by thresholding the experimental PLI by their median value so that all elements of the PLI matrix below the median value are set to zero (see Fig. 13).

The structural connectivity used to provide the network of the Hopf whole-brain network is the averaged dMRI connectome of the healthy control cohort (see Fig.1b). The structural connectivity is unitless and, hence, we use a scaling constant *C* as a free parameter. We found that the goodness of fit of the Hopf whole-brain model is not sensitive to structural connectivity scaling (see Fig. 8 in the Appendix). As long as the scaling parameter is large enough, it has no impact on the goodness of fit. As such, we parameterize the scaling at *C* = 20 and keep it this way for the remainder of the study. The natural frequencies of the individual brain regions must also be parameterized in the Hopf whole-brain model. We set the natural frequencies equal to the average peak frequencies of the experimental MEG signal of the healthy cohort. As such, the only remaining parameters of the Hopf whole-brain model are the global coupling strength *K* and the global excitability parameter *λ*.

We perform a two-dimensional grid search over the coupling strength and excitability parameter to find the optimal parameterization of the Hopf whole-brain model to average control phase-lag index connectivity as shown in Fig. 1d. Contrary to previous modeling assumptions, we find that the optimal fit to functional connectivity occurs at positive excitability parameter values (*λ >* 0). We also see that there are contour lines over the coupling strength and positive excitability parameters where the goodness of fit does not change. This suggests that the phase dynamics (as measured by PLI) are invariant—to some degree—under some transformation for positive excitability values.

By scaling the variables of the Hopf whole-brain model by 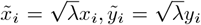 for *i* = 1 … *N*, we obtain the following equivalent system

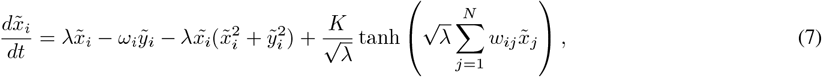

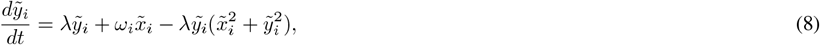

for *λ >* 0. The scaled system above is equivalent to a system coupling Hopf oscillators with a stable limit cycle of radius 1 with a modified coupling constant 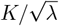. Additionally, the excitability parameter *λ* does not scale the radius of the limit cycle as in the original system. Instead, it is the timescale of the amplitude dynamics. As such, we expect the phase dynamics of the Hopf whole-brain model to be largely determined by the ratio 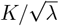 and not by the coupling strength and excitability independently. We see that this is indeed the case when computing the goodness of fit over coupling strength and excitability on a log-log grid (see Fig. 2) showing that the goodness of fit is invariant under 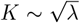. The coupling strength is a measure of interregional (global) coupling, whereas the excitability parameter is a measure of intraregional (local) excitability. As such, it is the competition between interregional and intraregional dynamics that determines the phase correlations in the Hopf whole-brain model. We will refer to the ratio 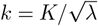 as the *normalized coupling strength*.

**Figure 2:**
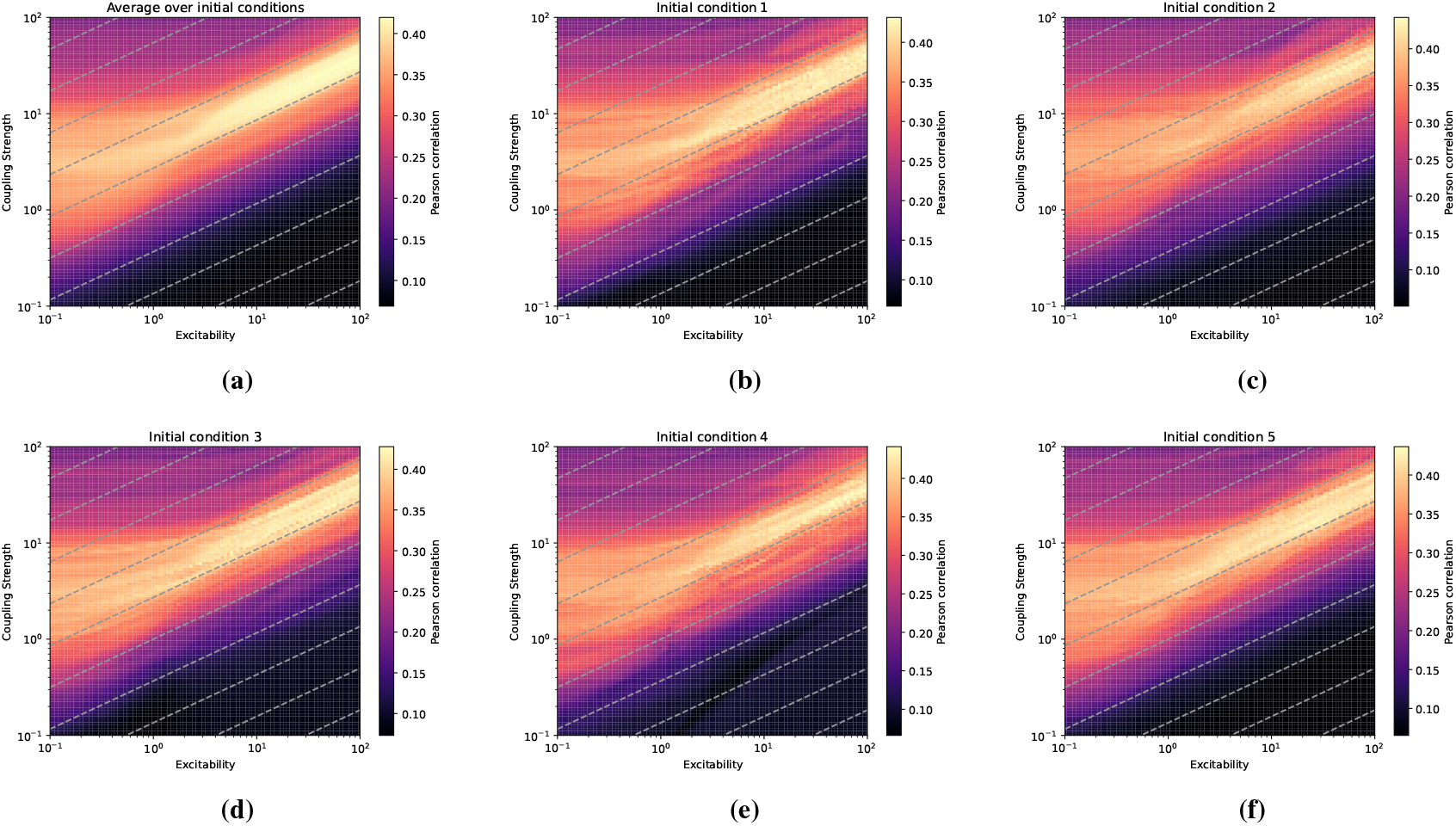
Parameter grid-search on logarithmic axes for a Hopf whole-brain model for individual initial conditions (b-f) and averaged over 300 initial conditions (a). The grey stippled lines represent a square-root scaling of the coupling strength with respect to excitability. That is, the grey stippled lines show 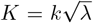 for different *k*. Each pixel shows the Pearson correlation between the simulated and experimental (average, control group) phase-lag index connectivity. The grids indicate that the Pearson correlation is unchanged as the coupling strength scales with the square root of excitability, such that 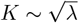.

### 3.2 Whole-brain modeling reveals higher interregional coupling in glioma cohort

We have now determined that it is the normalized coupling strength that controls the phase dynamics of the Hopf whole-brain model, and we can now investigate whether the normalized coupling strength differs between glioma patients and healthy controls. As before, we perform two-dimensional grid searches over the coupling and excitability parameters computing the goodness of fit to the averaged PLI of the healthy control group and the glioma group, respectively (Fig. 3b,c). We then find the optimal coupling strength per excitability parameter for both cohorts and compute the mean and standard deviation of the optimal coupling strength over randomized initial conditions for the whole-brain simulations (Fig. 3a, solid lines).

**Figure 3:**
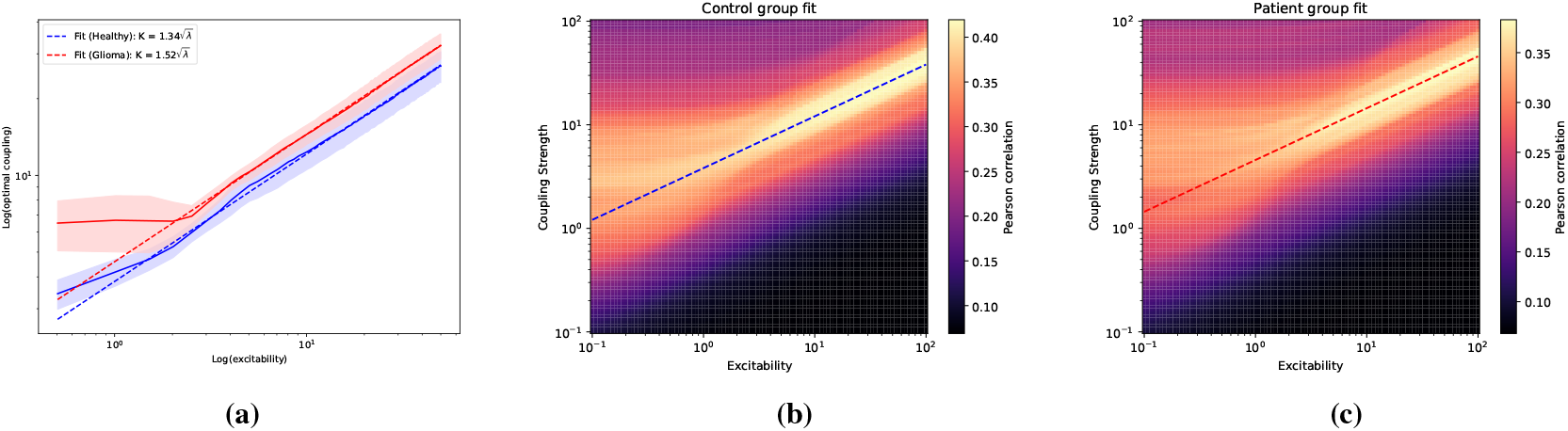
Optimal parameter fits of the Hopf whole-brain model to experimental phase-lag index connectivity for the control (blue) and glioma (red) cohort average. The optimal normalized coupling strength is fitted by regression (stippled line) and is the slope of the average optimal coupling strength per excitability parameter (solid line). Notice that all plots are in log-log coordinates. (a) The average (line) and standard deviation (shaded region) of optimal excitability and coupling strength over 1000 initial conditions. The average optimal coupling strength is fitted to a square-root dependence on the excitability (Hopf) parameter per the healthy (blue) and glioma (red) cohort. The regression fits the average optimal parameters well. (b,c) The fitted line (regression) of the optimal ratio between coupling strength and excitability parameters is shown on the parameter grid search for the healthy and glioma cohorts.

We find that the glioma cohort has a higher coupling strength over the excitability parameters, suggesting that their *normalized coupling strength* is higher. To confirm this, we find the optimal fit for the mean optimal coupling strength per the square root of the excitability parameter through least squares regression. That is, we find the scaling *k* providing the best fit for 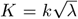 for the mean of the two cohorts. As shown in Fig. 3a, the regression provides an excellent fit for larger excitability values and produces a higher normalized coupling strength for the glioma cohort compared to the control cohort. The fitted regression lines of the normalized coupling strength have been plotted on top of the 2-dimensional grid searches in Figure 3b,c.

To further investigate the differences in functional connectivity between the control and glioma cohorts, we compute the distribution of optimal whole-brain parameters over the initial conditions for fixed excitability values. For 1000 initial conditions, we compute the difference in optimal coupling strengths between the glioma and control cohorts for five different excitability parameter values and plot the resulting distributions in Figure 4a-e. Interestingly, there is considerable variation between initial conditions. In particular, there seem to be two peaks in the distributions: one at zero difference and one at a positive difference. The bimodality disappears once all the optimal coupling strengths from the different excitability values are pooled into one distribution as shown in Figure 4f. Many initial conditions do not exhibit different optima between the glioma and control cohorts, however, those that do exhibit differences have a bias towards higher coupling strength in the glioma cohort.

**Figure 4:**
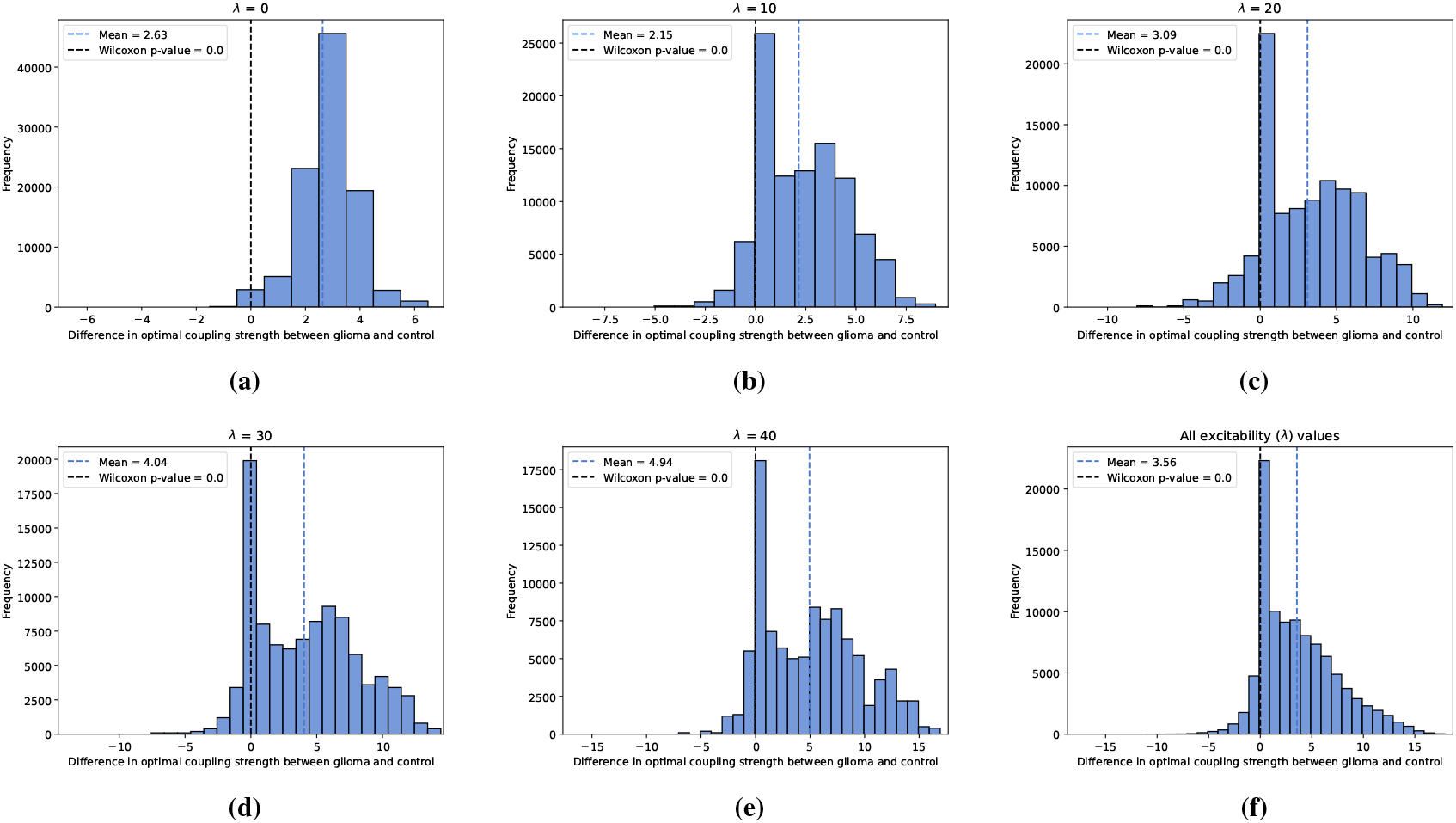
Distributions of the difference in optimal coupling strength between the healthy and glioma cohort for fixed excitability parameters over 1000 initial conditions for the Hopf whole-brain model. The black stippled line illustrates where there is zero difference between the cohorts, and the blue stippled line shows the mean of the distributions. (a-e) For different excitability (Hopf) parameter values, we find the difference in optimal coupling strength between the averaged control and glioma cohort when optimized for the best fit to experimental phase-lag index connectivity. The difference is positive when the glioma cohort has a higher optimal coupling strength, and negative when the glioma cohort has a lower optimal coupling strength. The distributions arise from 1000 simulations of the Hopf whole-brain model for random initial conditions. (f) In this histogram, all the optimal coupling strengths for all the excitability parameters (ranging from 0 to 50 with a step of 0.1) are pooled together into one larger distribution. For all distributions, the Wilcoxon rank-sum test was performed for which all were highly significant (to orders smaller than -100).

### 3.3 Tumors contribute to pathological interregional coupling in glioma patients

We have established that the normalized coupling strength (interregional versus intraregional dynamics) is higher in the glioma cohort average when compared to controls. However, we will see that individual patients also exhibit higher normalized coupling strengths. For fixed excitability values, we find the optimal coupling strength when fitting simulated PLI to the *patient-specific* PLI (thresholded by their median value). We then compute the difference between these optimal coupling strengths to that of the average control group over different initial conditions, as shown in Figure 5. As demonstrated in the previous section, considerable variations between initial conditions can exist. Nevertheless, we observe a general trend in which patients individually have higher coupling strengths than expected.

**Figure 5:**
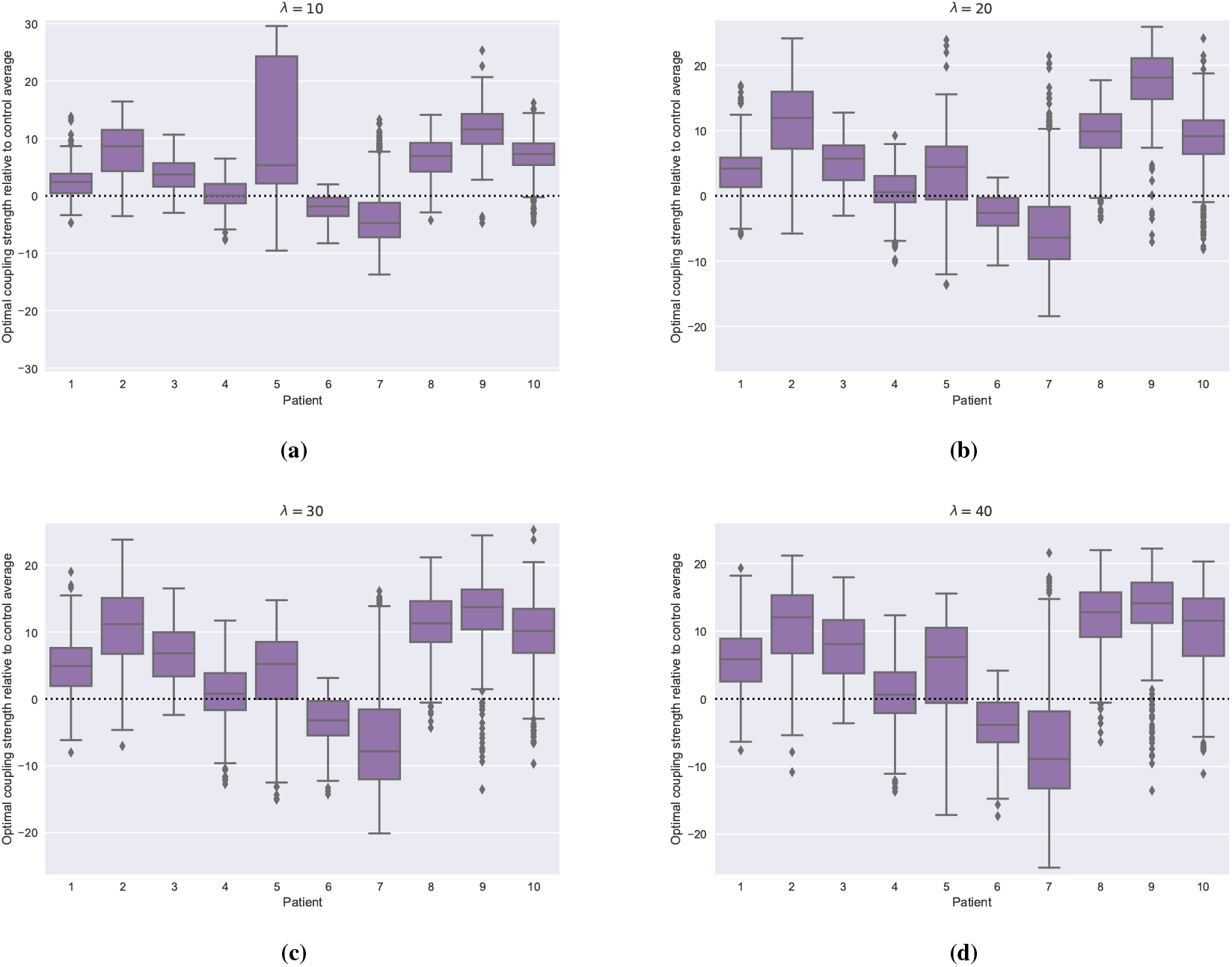
The difference in coupling strength found for individual patients and the healthy control, at different excitability parameter values *λ*. The box plots show the difference between optimal coupling strengths per patients’ individual phase-lag index compared to the optimal coupling strength found per the healthy control, averaged over 500 initial conditions for the Hopf whole- brain model. A box distributed towards positive values indicates that the patients’ optimal coupling strength is higher than the optimal coupling strength for the healthy control average.

To probe the role of tumors in shaping functional connectivity alterations, we perform a *pseudo-craniotomy* in which we remove the tumor regions of the particular patient from the goodness-of-fit evaluation when finding optimal model parameters. The regions themselves are not removed from the model as such a virtual procedure would be too disruptive, but the objective function (Pearson correlation between simulated and empirical functional connectivity) ignores them when identifying optimal parameters. As such, if the optimal coupling strength is lower after pseudo-craniotomy, it means that the tumors of that patient contribute to a pathologically high coupling strength.

The tumor regions of the patients in the glioma cohort are shown in Figure 6a-j with their position in the physical structural connectome. For each patient, we first find the optimal parameters with respect to their complete, individual PLI connectome. We then recompute the optimal parameters with respect to their individual PLI connectome after pseudo-craniotomy, that is, where the rows and columns of the tumor regions (all correlations involving the tumor regions) have been removed. As such, the change in optimal parameters reflects the impact the regions have on the goodness of fit of the whole-brain model and, by extension, their impact on the overall whole-brain dynamics.

**Figure 6:**
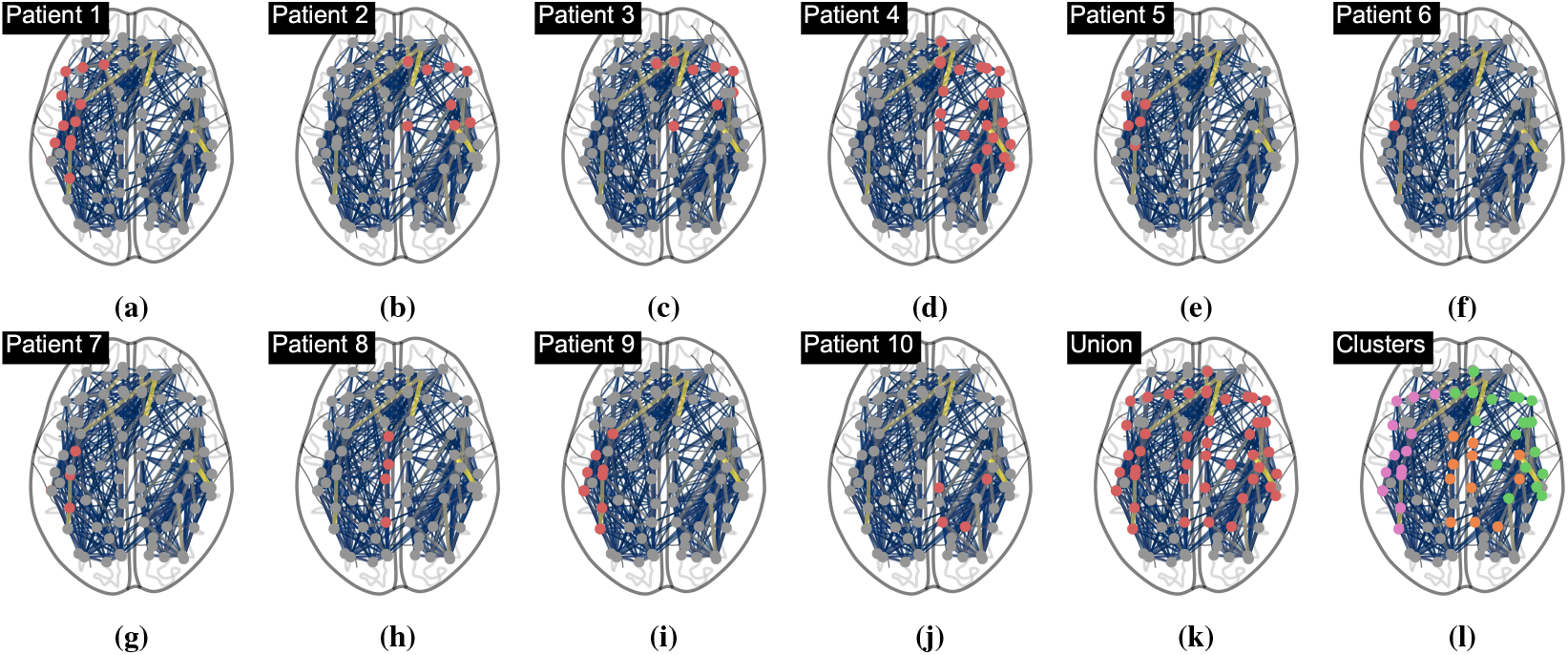
Pseudo-craniotomies were performed to assess the contribution of tumors to the whole-brain model dynamics. Here we show the locations of the tumor regions on top of the average structural connectivity. We also found three clusters (Kmeans clustering algorithm) of tumor regions which will be used in subsequent analyses. The union of all tumor regions will also be used.

We also identify three clusters of tumor regions by the K-means clustering algorithm, as shown in Figure 6k,f. In addition to investigating the tumor regions impact on global brain dynamics per patient, we also investigate the impact of the clusters—as well as the union of tumors across all patients—on the overall global dynamics, where we now fit the optimal parameters to the average PLI of the glioma cohort.

Setting the excitability parameter at *λ* = 40, we compute the optimal coupling strength before and after pseudo-craniotomy for each patient and tumor cluster (see Fig. 7). The goodness of fit is computed with patient-specific PLIs for each patient and averaged PLI (across patients) for each cluster. For each simulation, we perform the same patient-specific pseudo-craniotomy to the average healthy control (fitting to the average healthy PLI). To keep the control and patient fit as comparable as possible, we use the average healthy structural connectome for both control and patient-specific simulations. Simulations are repeated over 500 initial conditions with the resulting distributions shown in Figure 7(a,c). There is considerable variation between patients, with some showing a clear reduction in coupling strength, whereas others are difficult to distinguish from their respective control. In Figure 7(c,d), we show the distributions of the differences in the optimal coupling strength after pseudo-craniotomy for each patient and their control. Wilcoxon rank sum tests were computed for each patient/cluster/union showing highly significant results. Patients show either no change or a decrease in coupling strength after pseudo-craniotomy. We reiterate that decreases in coupling strength after pseudo-craniotomy correspond to the tumors contributing to higher coupling strengths.

**Figure 7:**
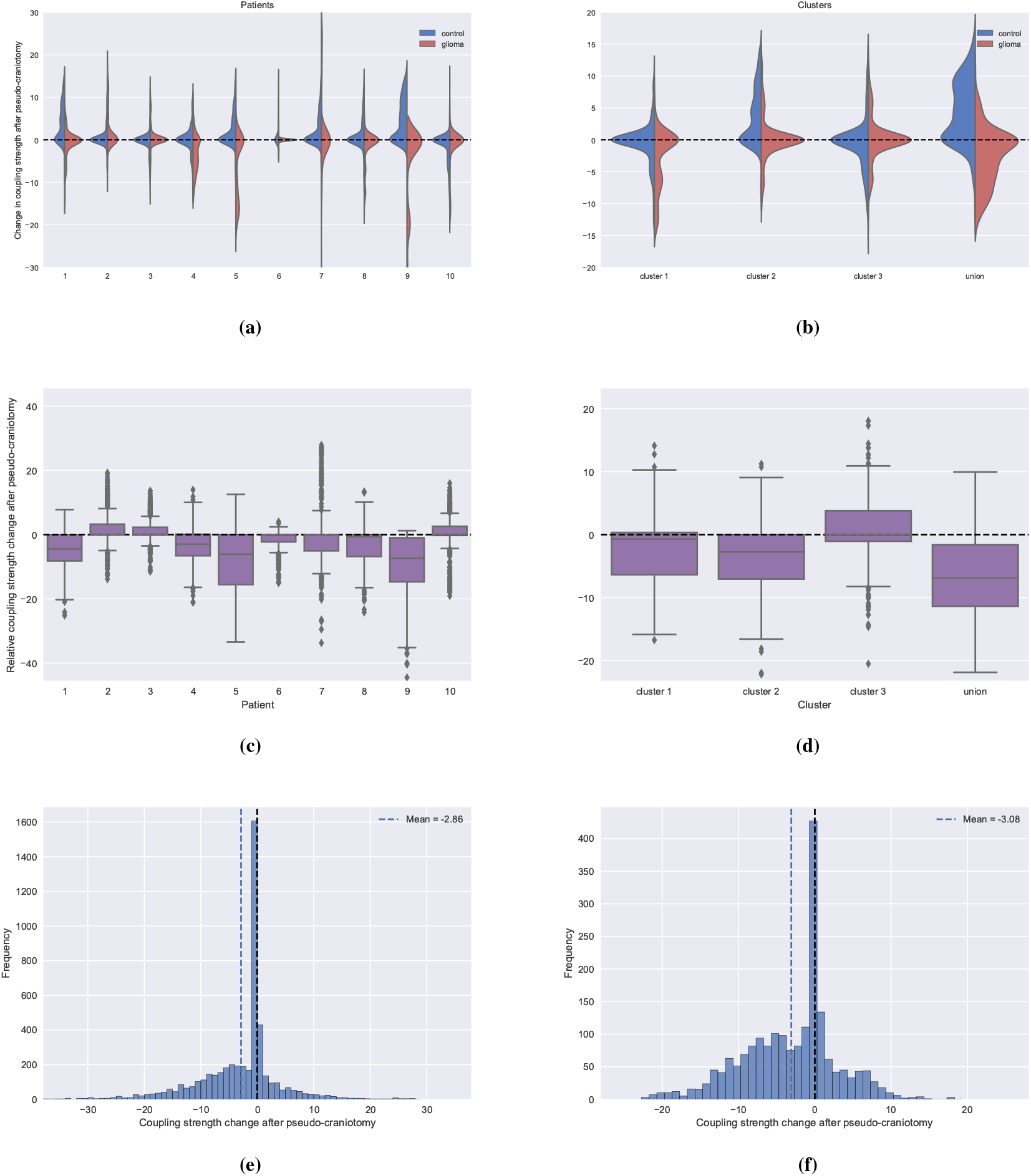
Pseudo-craniotomies were performed to assess the impact of tumors on the Hopf whole-brain dynamics. During pseudo- craniotomy, tumors are removed from the goodness-of-fit function changing the optimal parameter fit (but not removed from the structural connectome). Decreases in coupling strength after pseudo-craniotomy indicate that the tumors are contributing to higher coupling strengths. For all simulations, the excitability parameter was kept constant (*λ* = 40) for over 500 initial conditions for the Hopf whole-brain model. Simulations were performed per patient (left column) and per tumor cluster (right column). (a,b) Violin plots of the change in optimal coupling strength after pseudo-craniotomy. Optimal coupling strengths were found individually by computing the best fit to the patient’s phase-lag connectivity (red). For each, patient we performed a control with the same pseudo-craniotomy but fitted to the averaged healthy control (blue). (c,d) The difference in the change after pseudo-craniotomy between the patients and healthy average control. If the box plots are centered towards negative values, then there are larger reductions in optimal coupling strength for the patient than compared to controls. (e,f) A histogram showing the distribution of changes in optimal coupling strength across all patients and initial conditions relative to the change in the average healthy control.

**Figure 8:**
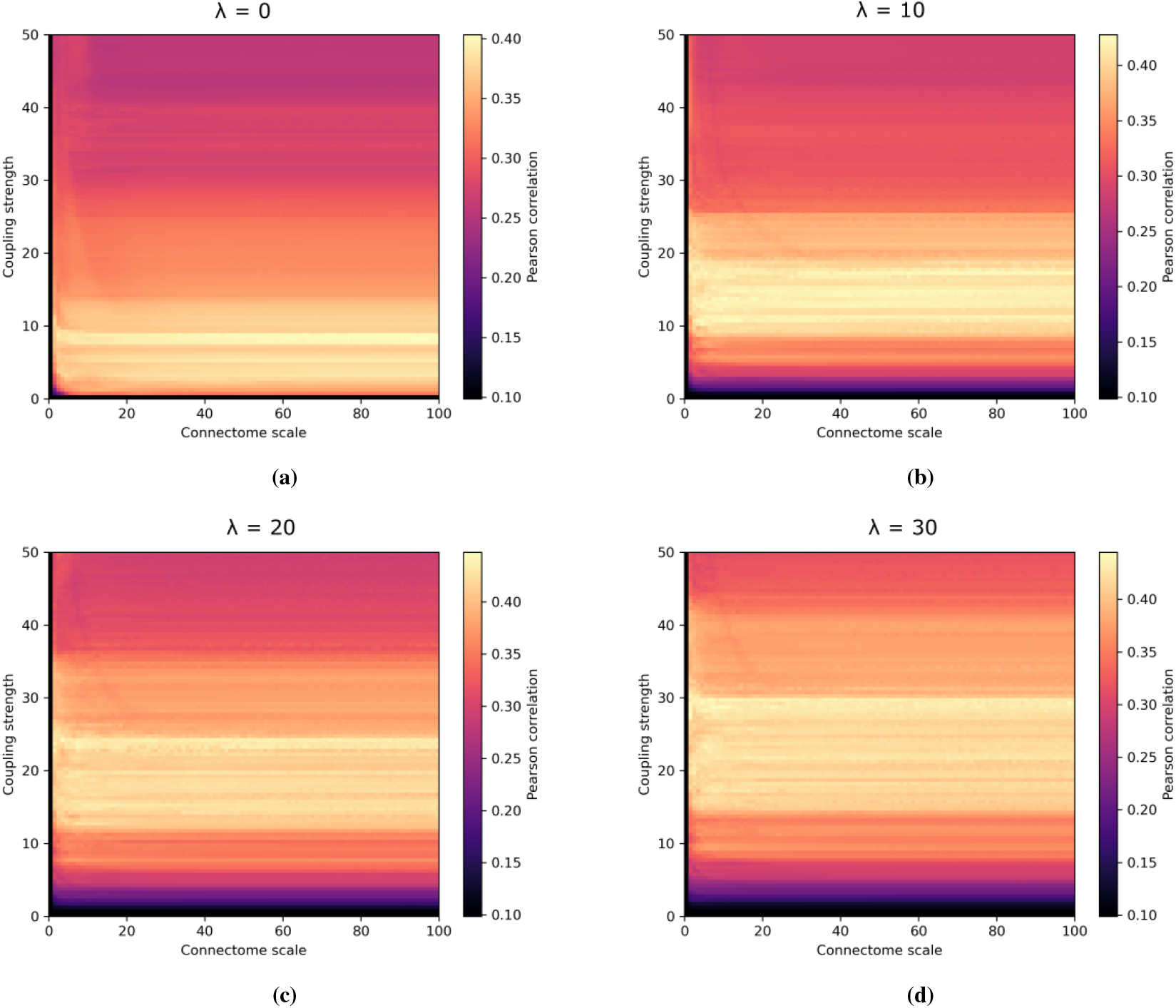
Grid search over coupling strength and connectome scaling for different excitability values for a set, random initial condition of the Hopf whole-brain model. Each pixel corresponds to the Pearson correlation between simulated and experimental (healthy control group) phase-lag index connectivity derived from MEG. As shown, the connectome scaling does not alter the model fit when sufficiently large, when varying the coupling strength and global excitability parameter *λ*. (a) *λ* = 0, (b) *λ* = 10, (c) *λ* = 20, (d) *λ* = 30

To further assess general changes caused by a pseudo-craniotomy, we pool the changes in optimal coupling (relative to the change in control) for the patients and clusters, as seen in Figure 7(e,f). For both patients and clusters, the distributions show a bias towards lower coupling strengths as induced by the pseudo-craniotomy. The Wilcoxon Rank Sum test was computed for the pooled patient and cluster distribution with a highly significant result, indicating a trend towards lower coupling strengths per the means of the distributions. Effectively, there is a general trend indicating that tumors contribute to higher coupling strengths, though, as expected, there is considerable variability between patients.

## 4 Discussion

We found that the optimal parameter fit of the Hopf whole-brain model to phase-based functional connectivity is invariant when 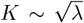 for *λ* ≥ 0. That is, the phase correlations of the simulated signal remain unchanged as the coupling strength scales with the square root of the excitability (Hopf) parameter. This is intuitive, as for isolated Hopf oscillators the radius of the stable limit cycle arising at the Hopf bifurcation (occurring at *λ* = 0) is given by 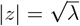. Mathematically speaking, 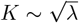 tells us that the phase dynamics does not change as the ratio between the coupling strength (interregional dynamics) and the radius of the self-sustained, local oscillations (intraregional dynamics) are constant. Interestingly, however, we find the optimal fit to functional connectivity at positive excitability parameters.

Contrary to common modeling assumptions [23, 51, 52], we did not find that the optimal parameter fit of the whole-brain model was close to criticality, in the sense that the local dynamics are *not* close to the Hopf bifurcation point(*λ* = 0). Indeed, we found the optimal fit to be arbitrary as long as the model was parameterized beyond criticality, with 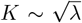 for *λ* ≥ 0. To be more precise, in our optimal fit, the individual local dynamics sustain oscillations independently of network interactions (the network interactions still affect the oscillations, but are not necessary to produce oscillatory signals locally). In most modeling studies, the parameters are set so that the individual oscillators can only sustain oscillations with the aid of network interactions. That is, the parameters are set close to a critical (bifurcation) point, which in our case would be *λ* ≤ 0. This assumption (*λ* ≤ 0) is based on the hypothesis that the brain is a dynamical system operating at criticality [53]. However, in our case, we have shown that the static, time-averaged functional connectivity—as measured by phase correlations—is better captured beyond criticality as defined by the Hopf bifurcation. Nonetheless, when considering the changes in phase correlation as a function of coupling strength, there is another sense of criticality as viewed through the lens of phase transition [54]. For example, for low coupling strengths, there will be no correlation patterns, however, patterns will arise as the coupling strength is increased, with an optimal coupling strength at intermediary values. In recent years, dynamic functional connectivity has found a growing interest which reveals the temporal evolution of functional connectivity. It seems likely that parameterizing the bifurcation parameter far beyond criticality *λ* ≫ 0 will *not* reproduce the switching between correlation patterns as observed experimentally with dynamic functional connectivity metrics [55].

Moreover, as revealed by scaling the Hopf whole-brain uniformly by 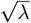, the excitability (Hopf) parameter acts as the time scale of the amplitude evolution for positive *λ*. As such, when *λ* is very high, the amplitude dynamics quickly adapt to the network interactions, whereas when *λ* is low, the amplitude dynamics is slow to adapt. As such, when *λ* is high, there is less fluctuation in the simulated signal amplitudes; the amplitude has little effect on the phase dynamics. Whereas, for small *λ* the amplitude experiences more fluctuations; the amplitude affects the phase dynamics. As such, we suspect the Hopf hole-brain model to behave similarly to a network of Kuramoto oscillators for high *λ* and that the square root scaling 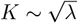 does not hold for small *λ* (the latter can be seen in Figure2). It is therefore reasonable to assume that amplitude-based metrics of functional connectivity will not be invariant under 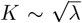. However, it is still unclear whether a positive or negative *λ* will reproduce more realistic amplitude correlations.

The Hopf whole-brain model is a phenomenological model. However, it has the advantage of using the Hopf normal form to describe the local (regional) dynamics. As such, close to criticality (Hopf bifurcation point at *λ* = 0), the Hopf normal describes the generic nonlinear behavior close to any Hopf bifurcation. Therefore, we can expect other whole-brain models employing more complicated biophysical neural mass models to behave similarly whenever they are parameterized close to a Hopf bifurcation point. However, we found the optimal fit of the Hopf whole-brain model to be arbitrarily beyond the Hopf bifurcation point. As such, it is not clear whether similar scaling laws between interregional (global) and intraregional (local) dynamics will exist in more complicated neural masses. It would be interesting to see whether a scaling law exists, for example, for the Wilson-Cowan model by performing a similar grid search for the coupling strength and a parameter known to traverse a Hopf bifurcation. In contrast to our study, more biophysical neural mass models have been used to provide insights into the whole-brain dynamics of glioma patients [35–37, 39].

Contrary to our results, studies by Aerts et al. [36] and Aerts et al. [37] do not find significantly different optimal coupling strengths between glioma patients and control subjects. However, the coupling parameter used in Aerts et al. [36] is more similar to the structural connectivity scaling parameter *C*, for which we do not expect any difference between cohorts, as we see that the structural scaling is underdetermined with respect to functional connectivity (see Fig. 8). In these studies, a more complex neural mass model—called the reduced Wong-Wang model—was used to fit fMRI functional connectivity. In addition to structural connectivity scaling, local inhibitory parameters were also tuned to guarantee biologically realistic firing rates. The local inhibitory strength was decreased for tumor regions and increased for nontumor regions. The exact analogy for the excitability parameter of the Hopf oscillator is not immediately clear in the reduced Wang-Wong model. However, if the overall effect of tuning the local inhibitory connections is to decrease the contribution of local dynamics to the emergent whole-brain dynamics, then we would expect the normalized coupling strength to increase as well. Similarly to the study by Aerts et al. [37], we find that the whole-brain model is quite robust to changes in dynamics after pseudo-craniotomy, though we find a general trend towards lower coupling strengths. That is, the relative contribution of the tumor regions to the distribution of optimal parameters is small, though existent. However, the optimal Hopf whole-brain model fit does not show the overall decrease in functional connectivity and changes in clustering and centrality as observed empirically in glioma patients (see Fig. 9).

**Figure 9:**
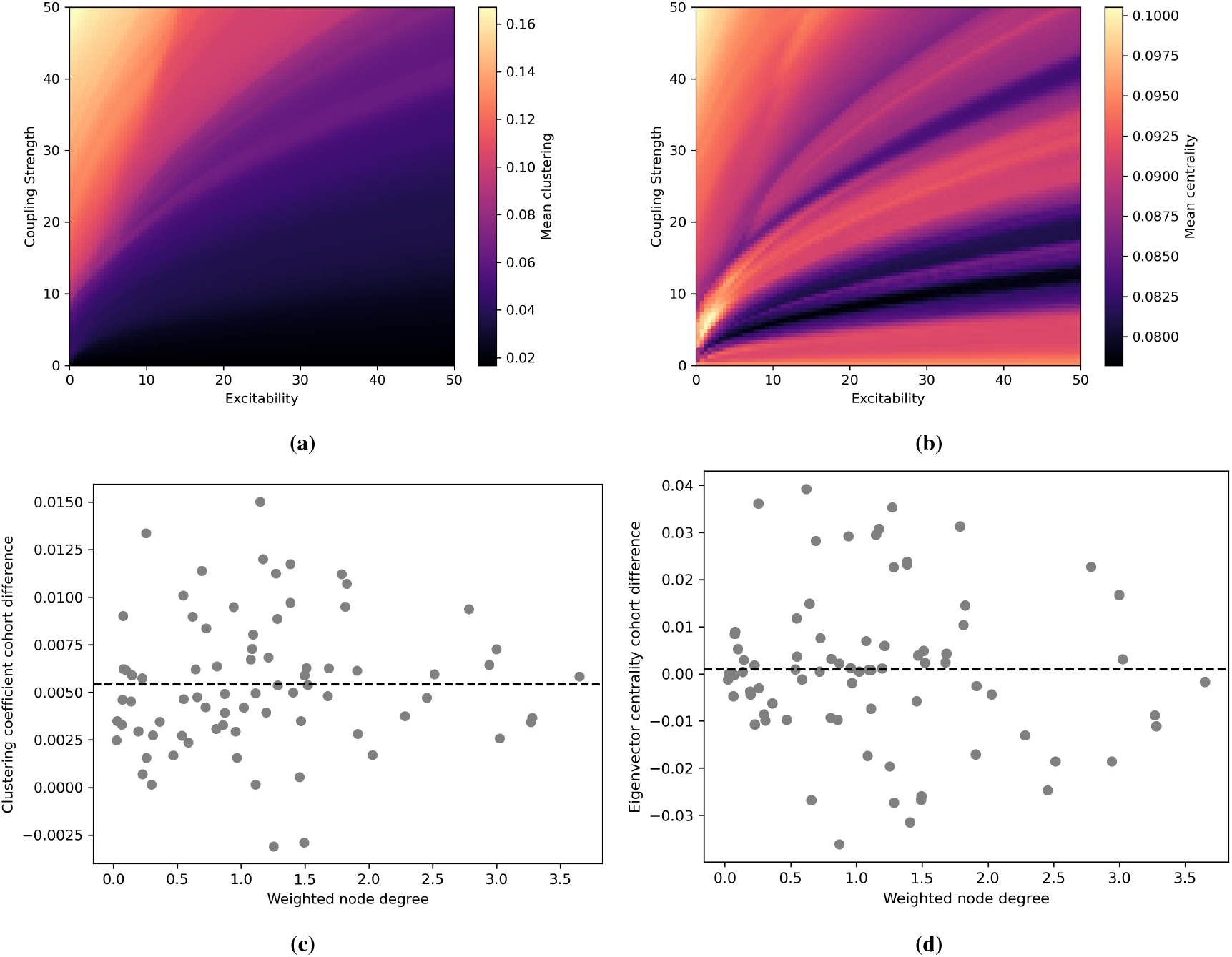
Network metrics of the simulated phase-lag index connectivity. (a) The average clustering coefficient of synthetic functional connectivity (phase-lag index) of the Hopf whole-brain model over the coupling strength and excitability parameter. The experimental clustering coefficient is around 0.2 for the healthy cohort. (b) The average eigenvector centrality over coupling strength and excitability. The empirical average eigenvector centrality is close to 0.05. (c,d) The difference in the simulated clustering coefficient and eigenvector centrality between the optimal model fit for the glioma and control group per node (plotted by their weighted structural node degree). The black stippled line shows the average difference, both of which are close to zero.

**Figure 10:**
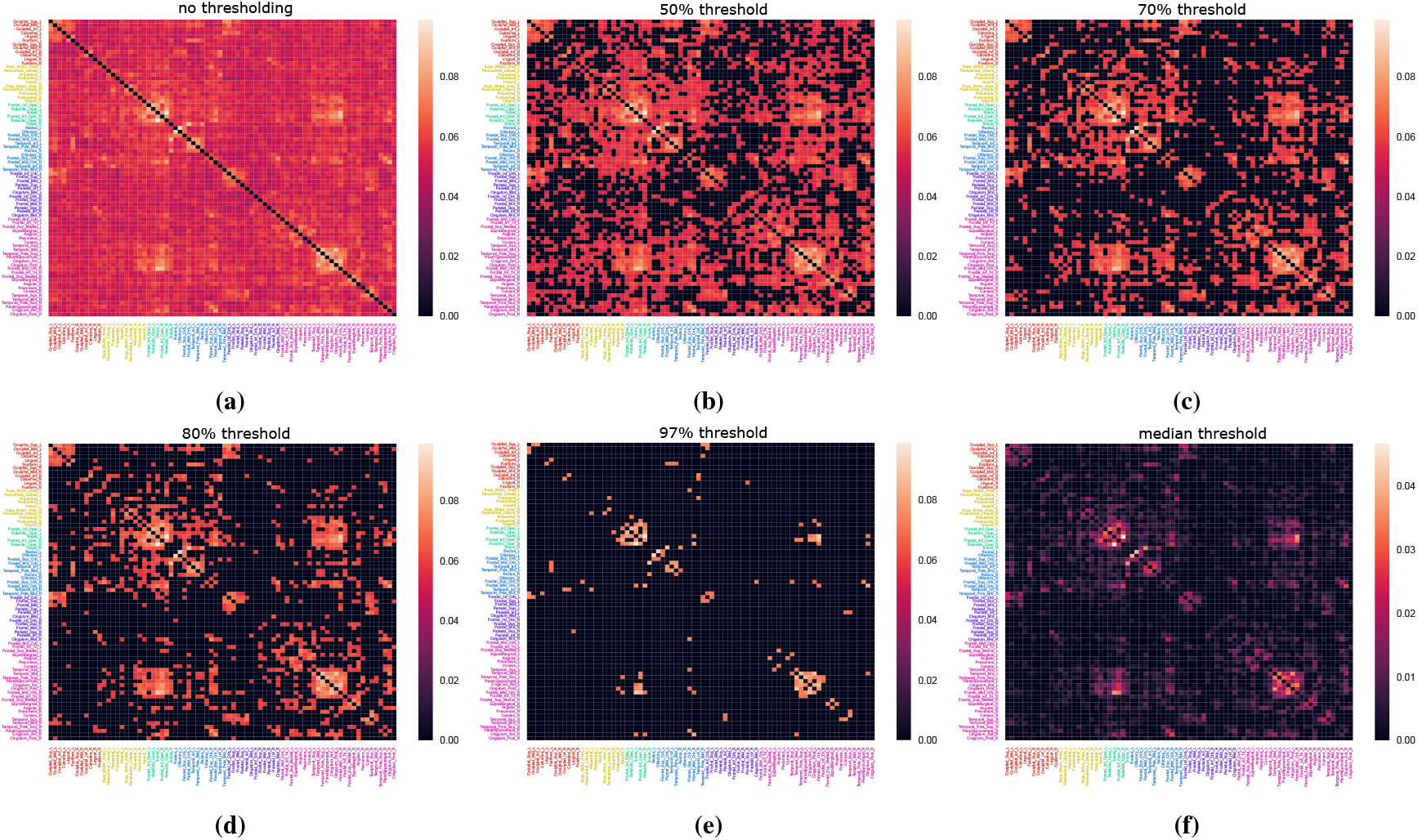
The post-processed average experimental phase-lag index connectome of the control group for varying levels of thresholding. During thresholding, all elements that belong to the lower X% are set to zero while others are kept at their original value. (a) before thresholding, (b) 50%, (c) 70%, (d) 80%, (e) 97% (which is the percentage giving the highest goodness of fit to simulated data), (f) Median thresholding; all values below the median of all elements are set to zero.

**Figure 11:**
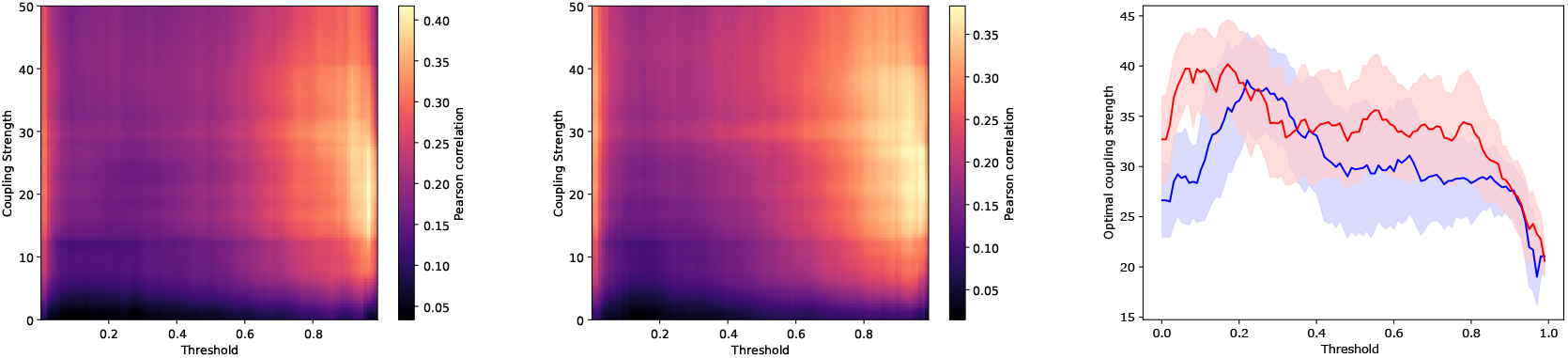
Grid searches showing the effect of coupling strength and thresholding of experimental phase-lag connectomes on the Pearson correlation between simulated and experimental phase-lag connectivity. The grid search matrices show the average of 300 initial conditions for the Hopf whole-brain model. (a) Grid search showing the fit to the average healthy phase-lag connectivity. (b) Grid search showing the fit to the average glioma patient phase-lag connectivity. (c) Plot showing the average (solid line) and standard deviation (shaded region) of the optimal coupling strength per experimental threshold for the control (blue) and glioma (red) cohort over initial conditions.

**Figure 12:**
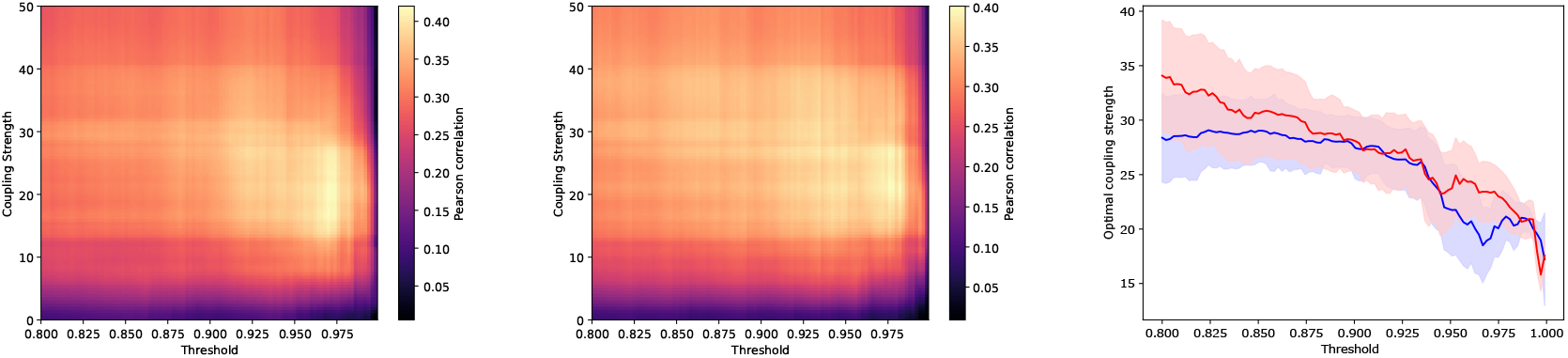
This plot is identical to Fig. 11 but zoomed in for higher threshold values.

**Figure 13:**
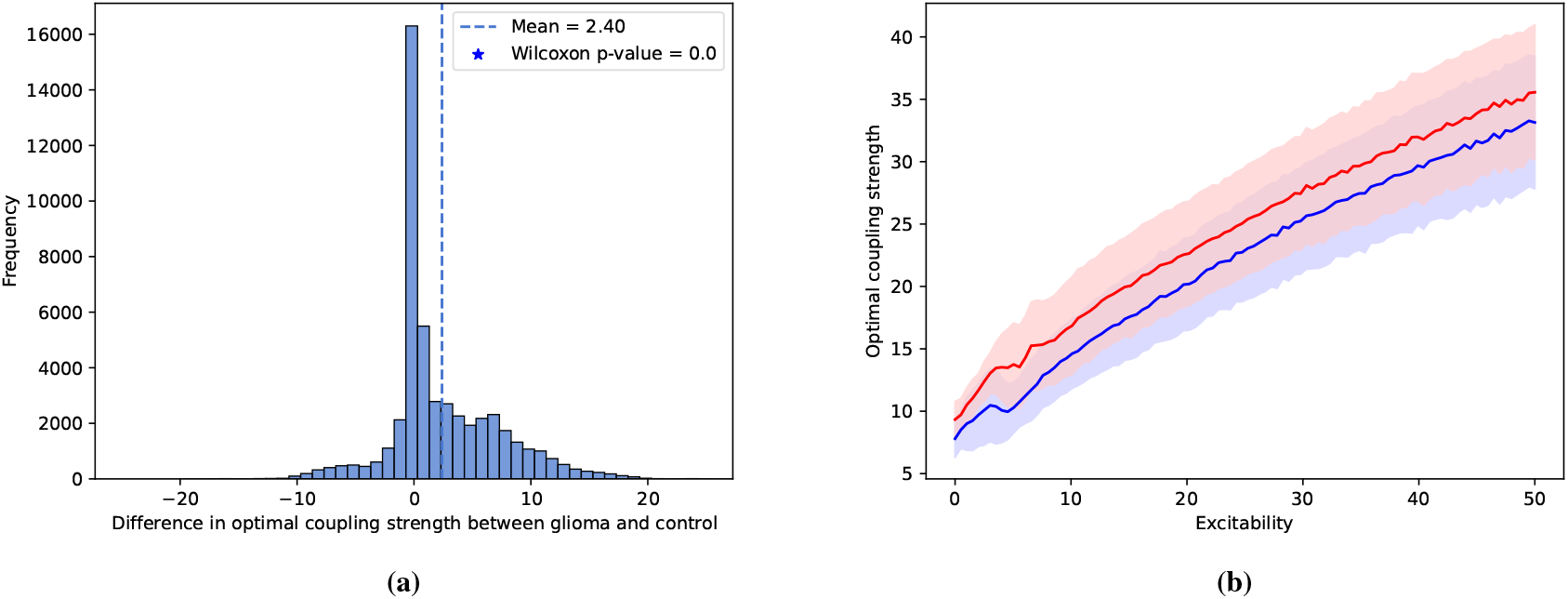
Optimal parameters for the Hopf whole-brain model fitted to PLI functional connectivity data when thresholded by their median value. The model was fitted to average PLI over each group for 100 randomized initial conditions. (a) The difference in optimal coupling strength (pooled over all excitability parameter values for 100 different initial conditions. (b) The mean (solid line) and standard deviation (shaded region) of optimal coupling strength per excitability parameter for the healthy (blue) and glioma cohort (red).

In conclusion, we showed that the phase dynamics of the Hopf whole-brain model is largely determined by the ratio of interregional and intraregional dynamics, where the ratio is shifted towards interregional dynamics in glioma patients More precisely, the phase dynamics is unchanged as the coupling strength grows with the square root of the excitability (Hopf) parameter. At the individual and population-based level, we find that glioma patients exhibit whole-brain dynamics with stronger interregional coupling than controls. Moreover, the tumor regions of individual patients contribute to this increase even though there is considerable variation across patients. In summary, we demonstrate an intimate link between global and local dynamics in whole-brain modeling and show that the contribution of global and local dynamics to emergent whole-brain dynamics is disrupted by the presence of tumors in glioma patients.

## Author contributions

CGA, LD, CB, and AG designed and conceptualized the study, discussed the results, and wrote the manuscript. CGA, CB, and AG developed the theoretical model and associated metrics. CGA performed all computations. MLMZ acquired and processed the data, and provided feedback on the results.

## Data Accessibility

Supporting data for this research will be given upon request.

## Ethics

The authors declare that they have no competing interests.

## Acknowledgements

We thank Lucas C. Breedt, Chris Vriend, and Marike van Lingen for their help with data acquisition and preprocessing.

## A Structural connectivity scaling

For varying levels of excitability values, we find that the goodness of fit of the Hopf whole-brain model is largely insensitive to the scaling of the structural connectome. As demonstrated in Figure 8, the scaling of the structural connectome just needs to be of sufficient magnitude to achieve optimal fit.

## B Network metrics of simulated functional connectivity

The network metrics of the simulated phase-lag index connectivity are not similar to the metrics found for empirical phase-lag index connectivity for optimal model parameters. There are only minor differences in network metrics between the optimal fit for the glioma and the control cohort, as shown in Fig. 9.

## C The effect of thresholding the empirical functional connectivity

The difference in optimal normalized coupling strength between the glioma and control cohort is for the most part positive across varying degrees of thresholding (see Figure 11 and 12). Both cohorts’ functional connectivity fit has its peak around 97% thresholding, which is the threshold used for the simulations in the Results unless specified otherwise.

When thresholding the glioma and control cohort by their median values (not the same thresholding value between the cohorts), we obtain similar results as for the optimal thresholding value (as shown in the Results) though with smaller differences in optimal coupling strength between the cohorts (see Figure 13).

